# Structural basis of aggregate binding by the AAA+ disaggregase ClpG

**DOI:** 10.1101/2023.08.31.555675

**Authors:** Panagiotis Katikaridis, Bernd Simon, Timo Jenne, Seongjoon Moon, Changhan Lee, Janosch Hennig, Axel Mogk

**Affiliations:** Center for Molecular Biology of Heidelberg University (ZMBH), DKFZ-ZMBH Alliance, Im Neuenheimer Feld 282, 69120 Heidelberg, Germany; German Cancer Research Center (DKFZ), Im Neuenheimer Feld 280, 69120 Heidelberg, Germany; Structural and Computational Biology Unit, European Molecular Biology Laboratory (EMBL) Heidelberg, Meyerhofstrasse 1, 69117 Heidelberg, Germany; Department of Molecular Biology and Biophysics, UConn Health, 263 Farmington Avenue, Farmington, CT, 06030-3305, USA; Department of Biological Sciences, Ajou University, Worldcup-ro 206, 16499 Suwon, South Korea; Chair of Biochemistry IV, Biophysical Chemistry, University of Bayreuth, Universitätsstrasse 30, 95447 Bayreuth, Germany

**Keywords:** ATPase associated with diverse cellular activities (AAA), protein aggregation, molecular chaperone, stress, 70 kilodalton heat shock protein (Hsp70)

## Abstract

Severe heat stress causes massive loss of essential proteins by aggregation necessitating a cellular activity that rescues aggregated proteins. This activity is executed by ATP-dependent, ring-forming, hexameric AAA+ disaggregases. Little is known about the recognition principles of stress-induced protein aggregates. How can disaggregases specifically target aggregated proteins while avoiding binding to soluble non-native proteins? Here, we determined by NMR spectroscopy the core structure of the aggregate targeting N1 domain of the bacterial AAA+ disaggregase ClpG, which confers extreme heat resistance to bacteria. N1 harbors a Zn^2+^-coordination site that is crucial for structural integrity and disaggregase functionality. Conserved hydrophobic N1 residues located on a β-strand were found crucial for aggregate targeting and disaggregation activity. Mixing experiments with N1-truncated AAA+ hexamers revealed that a minimal number of four N1 domains must be present in a AAA+ ring for high disaggregation activity. We suggest that multiple N1 domains increase substrate affinity through avidity effects. These findings define the recognition principle of a protein aggregate by a disaggregase, involving simultaneous contacts with multiple hydrophobic substrate patches located in close vicinity on an aggregate surface. This binding mode ensures selectivity for aggregated proteins while sparing soluble, non-native protein structures from disaggregase activity.

## Introduction

Bacterial Hsp100 chaperones constitute a subfamily of AAA+ proteins (ATPase associated with diverse cellular activities) and play crucial roles in proteostasis networks by acting on misfolded and aggregated proteins (1). They consist of one or two AAA domains, which mediate ATP binding and hydrolysis and oligomerization into hexameric rings. The energy derived from ATP hydrolysis is used to thread substrate proteins through an inner translocation channel. Substrate-binding pore residues of the AAA domains are arranged in a spiral staircase and propel the substrate in discrete steps that are orchestrated by cycles of sequential ATP hydrolysis events (2). While AAA+ proteins share this basic motor activity, they differ in cellular functions and act on diverse substrates. Functional specificity is gained by extra domains, which are either fused to or inserted into the AAA module (3, 4). These extra domains either directly interact with specific substrates or act as binding platforms for cooperating adaptor proteins, which deliver their bound cargo for subsequent processing by their AAA+ partners (5). Extra domains are typically connected to the AAA ring structure by flexible linkers, facilitating substrate binding and subsequent transfer to the processing pore site (6, 7). Extra domains can additionally regulate the ATPase activity of AAA+ proteins, allowing to adjust AAA+ motor activity to partner and substrate availability (8).

Hsp100 chaperones act on a variety of protein quality control substrates including soluble misfolded proteins, aggregated proteins and proteins harboring specific recognition sequences like the SsrA-tag. The SsrA-tag directly binds to the processing pore site of the Hsp100 members ClpA and ClpX (6, 9, 10). In contrast, ClpB/Hsp104 bind soluble unfolded proteins including the model substrate casein via an N-terminal domain (NTD) harboring a hydrophobic groove (11-13). The recognition of aggregated proteins by Hsp100s remains poorly characterized. Though ClpB/Hsp104 represent the canonical disaggregases of bacteria and fungi, they hardly bind to protein aggregates but require assistance by an Hsp70 partner chaperone for aggregate targeting (14, 15). Hsp70 binds first, coats the surface of protein aggregates and recruits ClpB/Hsp104 in a second step via physical interactions with their coiled-coil M-domains (16-19). ClpB/Hsp104 recruitment is directly coupled to ATPase activation, strongly enhancing their threading activities and enabling for disaggregation. Hsp70 binds both, soluble misfolded proteins including nascent polypeptide chains and aggregated proteins (20-22). Thus Hsp70 itself cannot discriminate between these two type of substrates. The selectivity of ClpB/Hsp104 is based on the number of Hsp70 molecules that have to interact with the AAA+ hexamer. A single Hsp70 is insufficient for ClpB/Hsp104 targeting, suggesting that a high local density of Hsp70 molecules serves as specific label for protein aggregates (16).

Bacterial ClpG (called ClpK in *Klebsiella pneumoniae*) represents a stand-alone disaggregase, which confers extreme heat resistance to bacteria (23, 24). ClpG harbors a distinct N-terminal domain (N1), which mediates protein aggregate targeting and is essential for ClpG activity (23, 25). ClpG has a high unfolding power demanding for tight control of its substrate specificity (26). How can ClpG distinguish between aggregated and soluble non-native proteins? Here, we dissected the selectivity of aggregate binding by ClpG by determining the structure of the N1 core domain via nuclear magnetic resonance spectroscopy (NMR). We identified hydrophobic N1 key residues crucial for aggregate targeting and show that multiple interactions between at least four N1 domains and the aggregate surface are required for disaggregation activity. These avidity effects enable ClpG to selectively target protein aggregates while sparing soluble, non-native proteins like nascent polypeptide chains from processing.

## Results

The ClpG N1 domain mediates the binding to aggregated proteins and can be transferred to ClpB, converting it from a partner-dependent to a stand-alone disaggregase (25). N1 thus functions as an autonomous unit. To understand the structural basis of N1 binding to protein aggregates we generated a structural model using AlphaFold2 (version AF2.2.2) (Fig. 1A). The model predicts an N-terminal (M1-V46, high confidence with pLDDT values >90%, Fig. S1A/B) core structure that is followed by an α-helical segment (S47-G71, low confidence with pLDDT values < 60%). The core harbors a putative Zn^2+^-binding center formed by the highly conserved residues C6/C9/C31/H34 (Fig. 1A, Fig. S1C). To determine the minimal part of N1 needed for aggregate targeting, we generated two ClpG deletion mutants lacking either the N-terminal core structure (ClpG-Δ2-46) or the C-terminal α-helix (ClpG-Δ47-71) and tested for refolding of aggregated Luciferase (Fig. 1B). As reference we used ClpG-ΔN1 (Δ1-106) lacking the entire N1 domain and a spacer sequence linking N1 to N2. Disaggregation activity of ClpG-Δ2-46 was strongly reduced and similar to ClpG-ΔN1, while ClpG-Δ47-71 was more potent than ClpG-WT. ATPase activities of all N1 deletion constructs were increased compared to ClpG-WT (Fig. 1C) and highest for ClpG-Δ47-71. This excludes that loss of ClpG-Δ2-46 disaggregation activity is caused by impaired ATP hydrolysis. Furthermore, the enhanced ATPase activities of all N1 deletion mutants point to an additional regulatory function of the N1 domain. The increased ATPase activity of ClpG-Δ47-71 correlates with its higher disaggregation activity.

**Figure 1.**
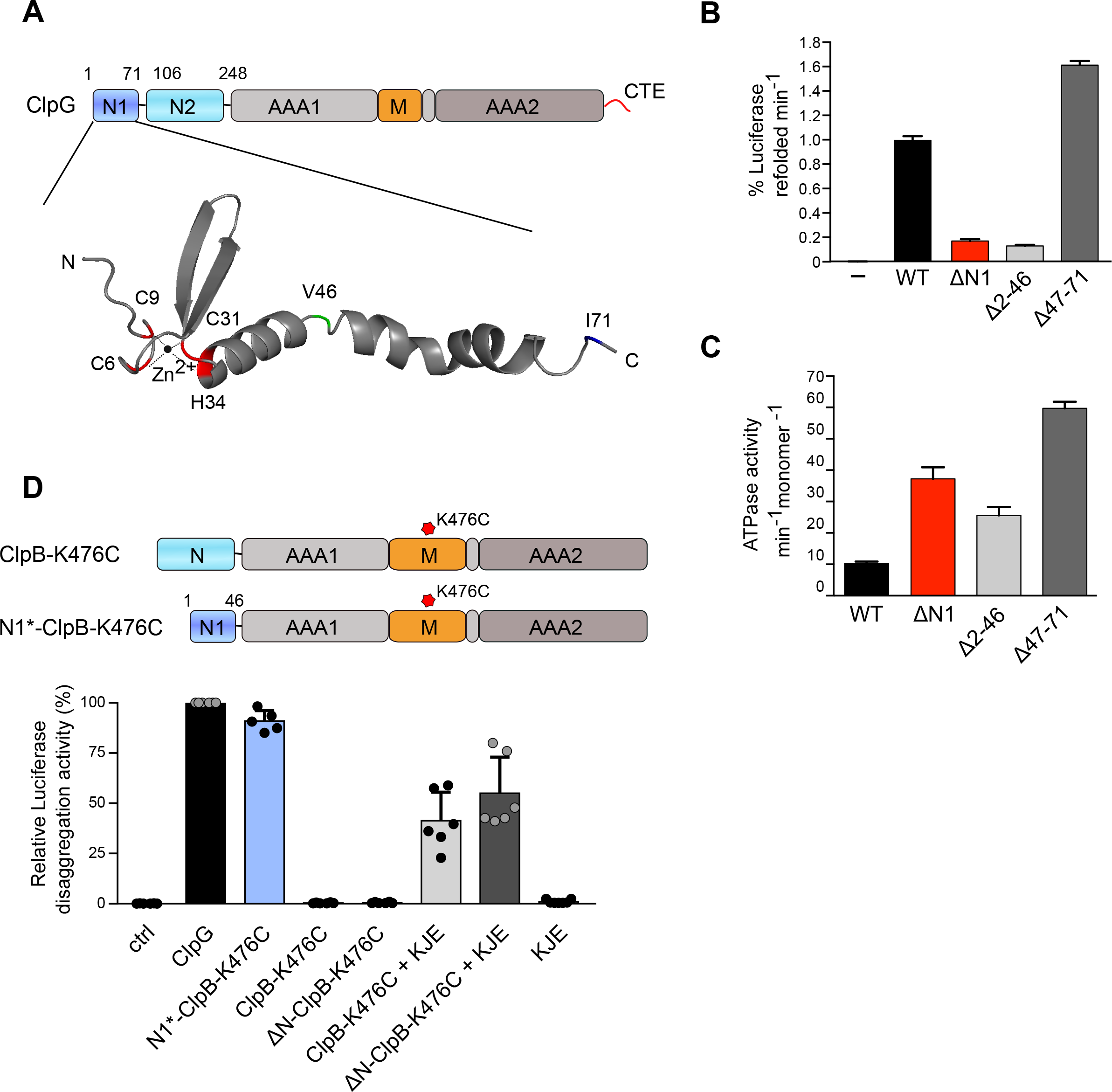
ClpG 1-46 (N1 *) is essential and sufficient for aggregate targeting. (A) Domain organization of ClpG. ClpG consists of two N-terminal domains (N1, N2), two AAA domains (AAAl, AAA2), a middle domain (M) and a C-terminal extension (CTE). Domain boundaries of N1 and N2 are indicated. An AlphaFold2 model of N1 is provided. Conse1ved residues implicated in zn^2+^ coordination are highlighted in red. (B) Luciferase disaggregation activities of ClpG WT and indicated N1 deletion constructs were determined. A control reaction lacking ClpG is provided. (C) ATPase activities of ClpG WT and indicated N1 deletion constructs were determined. (D) Domain organizations of ClpG-ClpB chimeras. The ClpB M-domain mutation K476C abrogates repression of the ClpB ATPase motor. Disaggregation of aggregated Luciferase by ClpG_N1-ClpB-K476, ClpB-K476C, N-ClpB-K476 (with and without the DnaK chaperone system (KJE)) was monitored by dete1mining Luciferase activities. The disaggregation activity of ClpG was set to 100%. Standard deviations are based on at least three independent experiments (B-D).

To provide direct evidence that the N1 core domain (N1*) is sufficient for aggregate targeting, we fused the N-terminal 46 N1 residues to ΔN-ClpB-K476C lacking its N-terminal domain. The mutation K476C resides in the ClpB middle-domain and overrides ClpB activity control allowing for high ATPase activities in absence of the Hsp70 partner (27). N1*-ClpB-K476C had similar ATPase activity as ClpB-K476C and ΔN-ClpB-K476C (Fig. S1D). However, the fusion construct showed high disaggregation activity while controls ClpB-K476C and ΔN-ClpB-K476C required presence of the Hsp70 partner for aggregate targeting and thus disaggregation activity (Fig. 1D). This demonstrates that N1* represents the aggregate targeting unit of ClpG that can be transferred to other AAA+ family members.

### Solution structure determination of the N1 core domain and identification of substrate binding residues by NMR spectroscopy

We aimed at determining the structure of the N1 core domain (N1*) to validate the AlphaFold2 prediction and at identifying residues involved in substrate binding by NMR. Purified N1* eluted as monomer in Superdex S30 size exclusion chromatography runs (Fig. S2A) making it a suitable target for NMR analysis. A longer N1 construct involving residues 1-82 also eluted as monomer, excluding a function of the N1 α-helical C-terminal part in dimer formation (Fig. S2A).

NMR chemical shift assignments and structure determination were performed by standard methods (see experimental procedures). The ^1^H,^15^N-HSQC spectrum showed well-dispersed peaks of which all could be assigned unambiguously (Fig. 2A). TALOS (28) derived secondary structure predicted from secondary chemical shifts confirms the presence of a β-sheet followed by an α-helix (Fig. S2B). The α-helix ends with Q41 and TALOS predicts that residues 42-46 are flexible. NOE assignments were difficult due to peaks exhibiting line broadening in the sequence range towards the tip of the β-sheet. The line broadening also results in a smaller number of NOEs per residue in this region, resulting in larger backbone rmsd values in the final ensemble of structures (Fig. 2B, see Table 1 for NMR structure calculation statistics). The initial structure calculation without an explicit Zn^2+^ ion exhibited a well-defined orientation of the side chains of C6, C9, C31 and H34 indicating the coordination of a Zn^2+^ ion, with the histidine side chain NE2 atom close to a potential Zn^2+^. The final calculation includes the Zn^2+^ ion coordinated in this orientation (29) (Fig. 2B). The mean structure superimposes well with the structure predicted by AlphaFold2 (Fig. S2C). However, the ensemble indicates dynamics that AlphaFold2 cannot predict. The structure is well defined until Q41 where the α-helix ends and the backbone conformation of the last five residues is random. The lack of structure in this region is confirmed by ^15^N relaxation data (Fig. S2). Heteronuclear NOE values drop below 0.7 from Q42 to the C-terminus (Fig. S2D) and model-free Lipari Szabo order parameters S^2^ are gradually reduced towards the end of the α-helix (Fig. S2E). The transverse relaxation rates *R*_*2*_ of V17, T27 and L38 are significantly increased and indicate break-points in the structure leading to slow exchange motions of residues 17-27 and 38-41 (Fig. S2F). The residues surrounding the Zn^2+^ ion exhibit local dynamics on a time scale that is faster than the overall tumbling rate (4.2 ns), except for V8, which is exchange broadened and thus moves at a slower time scale. The ^15^N relaxation data support the result of the structure calculation of a well-defined molecule around the Zn^2+^ ion and increased structural heterogeneity of residues V17-T27 and L38-Q42.

**Table 1.**
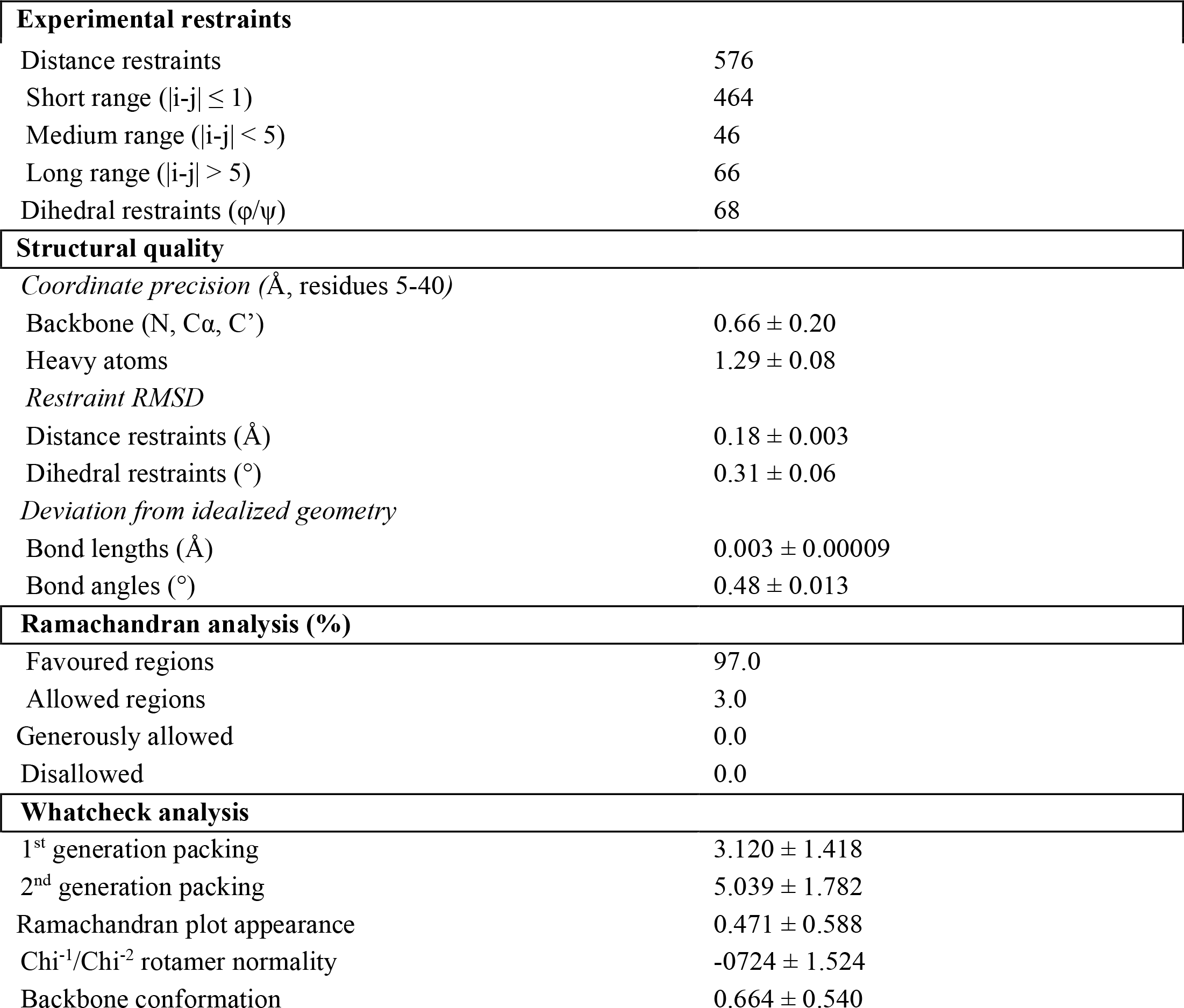
Structural statistics of the N1* domain NMR ensemble (structural validation was performed using CING (46) inside NMRbox (47)

**Figure 2.**
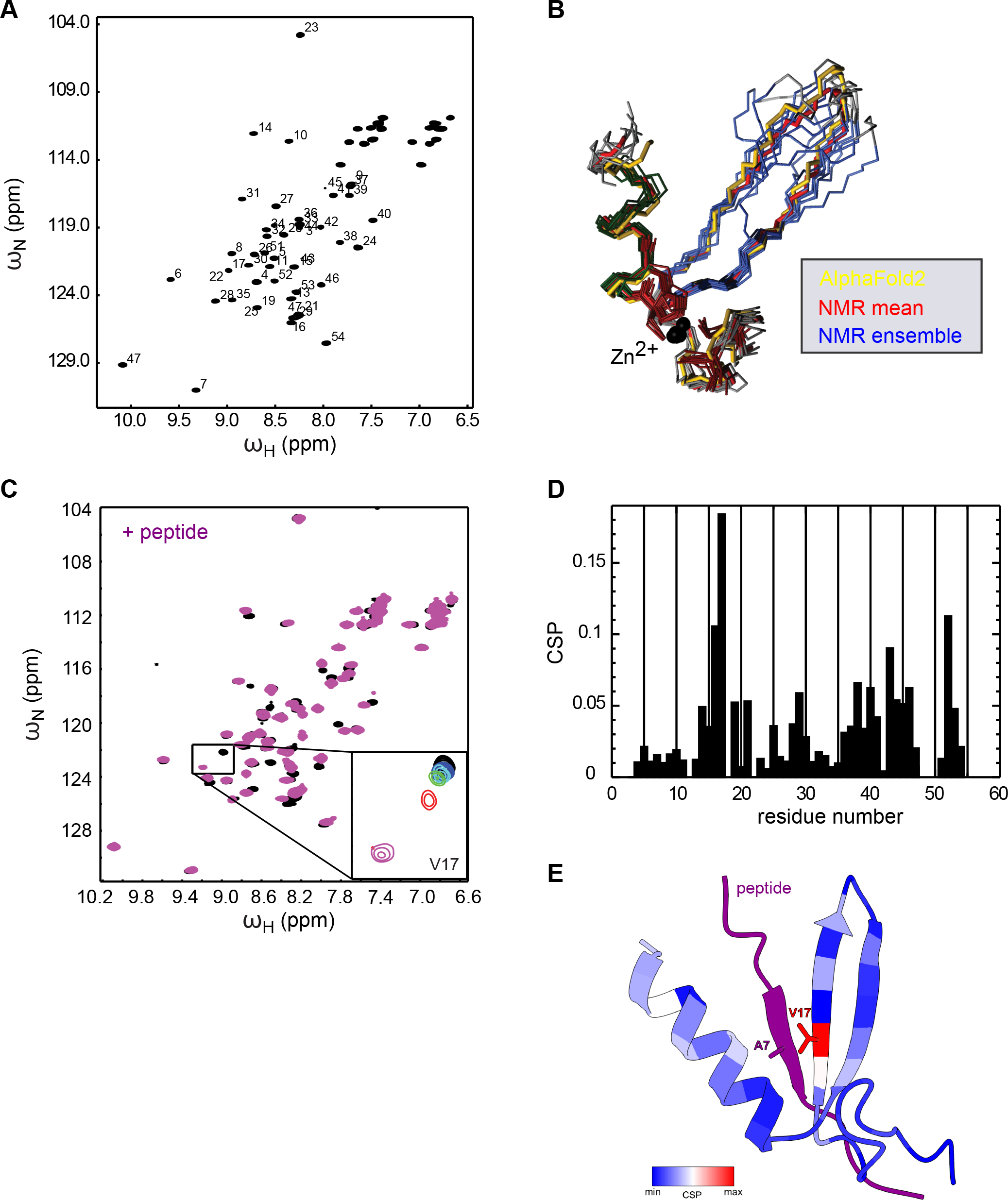
N1 * NMR structure determination and substrate binding site mapping by NMR spectroscopy. (A) ^1^H-^15^N HSQC spectrum of the N1 *domain.Numbers at cross peaks are the residue numbers assigned to the resonances derived from backbone assigmnent experiments. (B) N1 * NMR structure ensemble (blue, ten lowest energy structures) superimposed with the structure closest to the mean (red) and the AlphaFold2 prediction (yellow). The Zn^2+^ ions are shown as black spheres.(C) Overlay of ^1^H-^15^N HSQC spectra of free N1 * domain (black) and after addition of 3-fold excess of peptide 134 (purple, 80 μM N1 *: 240 μM peptide 134). The strongest CSP is exhibited by Vl 7 and shown in a zoom view as inset in the full spectrum with four more inte1mediate titration steps. (D) Chemical shift pe1turbation magnitude plotted versus the amino acid sequence of the N1 * domain. (E) AlphaFold-Multimer structure prediction of the N1 *-peptide 134 complex. The CSPs values from (D) are plotted onto the structure with blue to red for weak to strong CSPs.

Peptide 134 (CWFLRYLKASKVPLN) was previously identified as N1 binder in a peptide library screen using *S. cerevisiae* Has1 as artificial model protein (25). Peptide 134 has a positive net charge and is enriched for hydrophobic and aromatic residues. To verify the interaction of Peptide 134 and N1*, we titrated the unlabeled peptide to the ^15^N-labeled N1* domain and monitored the chemical shift perturbations of the backbone HN moieties in ^1^H,^15^N-HSQC spectra (Fig. 2C/D). All perturbed resonances are gradually changing frequencies and are thus in fast exchange indicating a binding affinity in the micromolar range. Several resonances exhibit additional line broadening upon peptide titration, including V17 for which the strongest chemical shift perturbation is observed (Fig. 2D). Based on this, the binding site of the peptide can be mapped to the β-sheet around V17, to the end of the C-terminal helix and the following unstructured residues (Fig. 2D). The chemical shift perturbations are not saturated at a 1:3 ratio of protein to peptide and at higher ratios the sample quality started to deteriorate. Therefore NMR data do not allow an exact fitting of a dissociation constant but can be estimated to be larger than 50 μM. We also predicted the N1*-Peptide 134 complex structure using AlphaFold2 (v2.2.2, Fig. 2E, S2G). Here, the peptide interacts with the N1* β-sheet forming an additional anti-parallel β-strand. The central residues of the β-completion are V17 of N1* and A7 of Peptide 134. The ?-terminus of the bound peptide is close to the C-terminal helix of N1*. In conclusion, chemical shift perturbations largely validate the AlphaFold2 model of the complex. More detailed NMR analysis of the structure and dynamics of the complex were hampered by insufficient long-term stability of the complex at higher concentrations.

### Zn^2+^-binding to N1* is crucial for structural integrity and function

The NMR structure strongly suggests that N1* binds Zn^2+^ via the conserved residues C6/C9/C31/H34. We directly show Zn^2+^ binding to ClpG-WT by ICP-EOS (Inductively coupled plasma optical emission spectrometry) measurements (Fig. 3A). Zn^2+^-binding to ClpG-ΔN1 (1-106) and ClpG-C6A/H34A was strongly reduced, indicating metal ion coordination via N1. Indeed, isolated N1* bound Zn^2+^ with the same efficiency as ClpG-WT (Fig. 3A). Comparison of isolated N1* with N1*-C6A/H34A by circular dichroism (CD) and NMR spectroscopy revealed strong differences indicative of protein unfolding (Fig. S3A/B). Loss of Zn^2+^-coordination in ClpG-C6A/H34A strongly increased basal ATPase activities, confirming the function of N1 as regulator of ClpG ATPase activity (Fig. S3C). ClpG-C6A/H34A did neither exhibit disaggregation activity *in vitro* nor *in vivo* and, accordingly, did not restore thermotolerance in *E. coli ΔclpB* mutant cells lacking the canonical ClpB disaggregase (Fig. 3B-D). ClpG-C6A/H34A was produced at similar levels as ClpG-WT documenting specificity of activity loss (Fig. S3D/E). We also tested ClpG-C6A/H34A activity in the authentic organism *Pseudomonas aeruginosa* (*Pa*), showing that its ability to restore heat resistance is completely lost in a *Pa* Δ*clpB* Δ*clpG* Δ*clpG*_*GI*_ mutant, which lack disaggregation activity (Fig. S3F/G). Together these findings demonstrate that Zn^2+^-binding is crucial for structural integrity of N1 and ClpG disaggregation activity.

**Figure 3.**
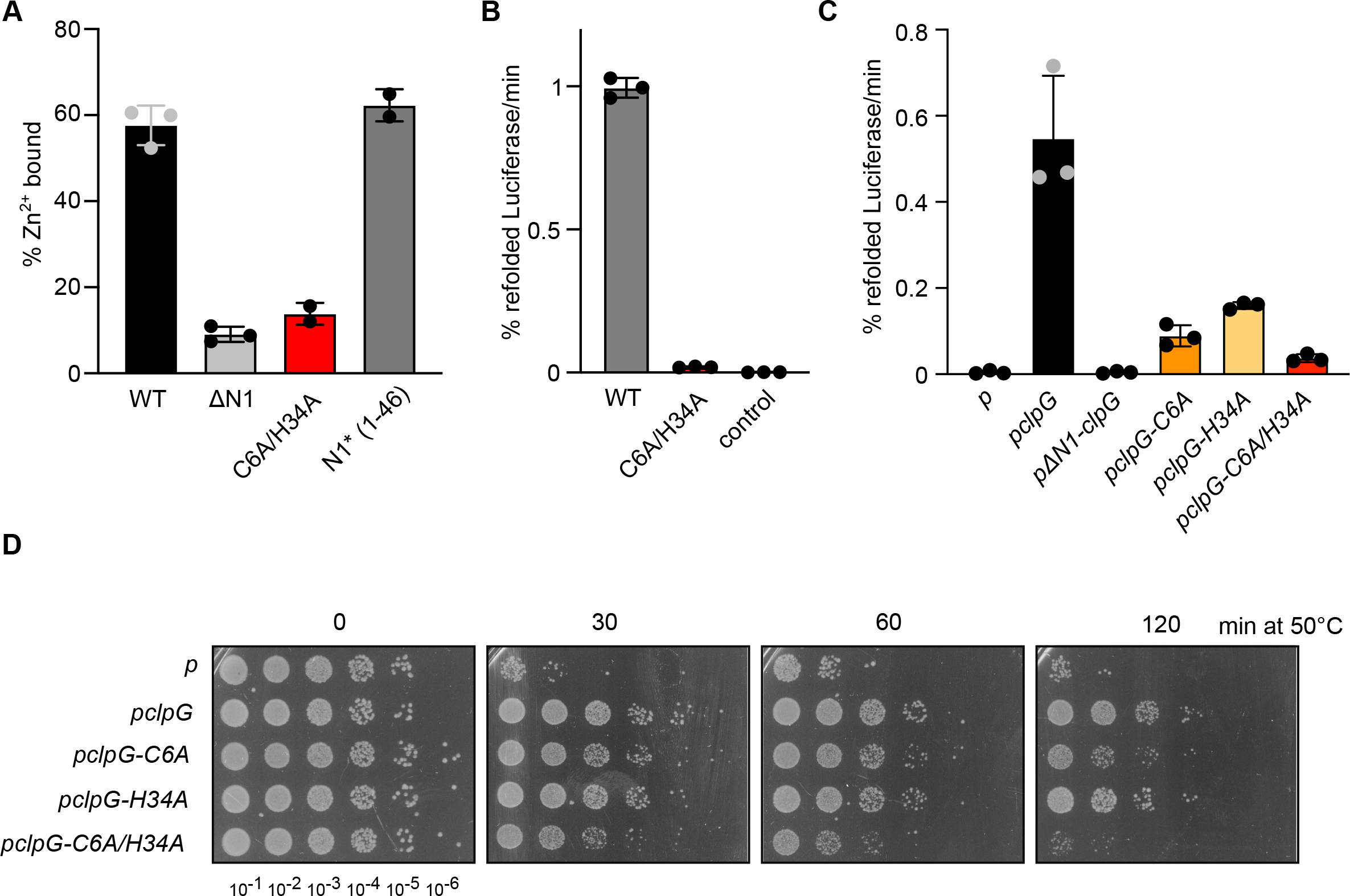
zn^2+^-binding is essential for ClpG disaggregation activity. (A) Zn^2+^-binding to ClpG, indicated mutants and isolated N1 was dete1mined by ICP-OES. (B) Luciferase disaggregation activities of ClpG WT and Zn^2+^-binding deficient ClpG-C6A/H34A were dete1mined. A control reaction lacking ClpG is provided. (C) *E. coli 1’1clpB* cells harboring plasmids for constitutive expression of Luciferase and IPTG controlled expression of *clpG* (wt and indicated mutants; p: empty vector control) were grown at 30°C to mid-logarithmic growth phase. Cells were shifted to 46°C for 15 min and Luciferase activities were determined during a recove1y phase at 30°C. (D) *E. coli 1’1clpB* cells harboring plasmids for expression of *clpG* (wt and indicated mutants; p: empty vector control) were grown at 30°C to mid-logarithmic growth phase and shifted to 50°C. Serial dilutions of cells were prepared at the indicated time points, spotted on LB plates and incubated at 30°C. Standard deviations are based on at least three independent experiments (A-C).

### Validation of the N1 substrate binding site

To ultimately determine and validate the N1 substrate binding site we mutated conserved N1 residues that showed strongest chemical shift perturbations in presence of Peptide 134: V17, L21, Q37, L38, Q41 and K43 (Fig. 4A). We generated respective glutamate mutants in a fluorescent ClpG-YFP fusion construct and expressed the constructs in *E. coli ΔclpB* mutant cells. This approach enabled us to determine *in vivo* disaggregation activities of ClpG mutants and to directly link alterations in activity status to changes in aggregate targeting. While the *in vivo* disaggregation activities of most N1 mutants hardly differed from ClpG-WT, reduced activities were determined for ClpG-L21E-YFP and ClpG-V17E-YFP (Fig. 4B). Notably, both residues are part of the first β-strand, whereas none of the mutations located in the α-helix showed reduced activity. Disaggregation activity was further reduced upon combining V17E and L21E mutations in ClpG-V17E/L21E-YFP, whose activity was similar to the Zn^2+^-binding mutant ClpG-C6A/H34A-YFP (Fig. 4B). All YFP-fusion constructs were expressed at comparable levels, except for ΔN1-ClpG-YFP, which was expressed at lower levels even when gene expression was fully induced upon addition of 1 mM IPTG (Fig. S4A).

**Figure 4.**
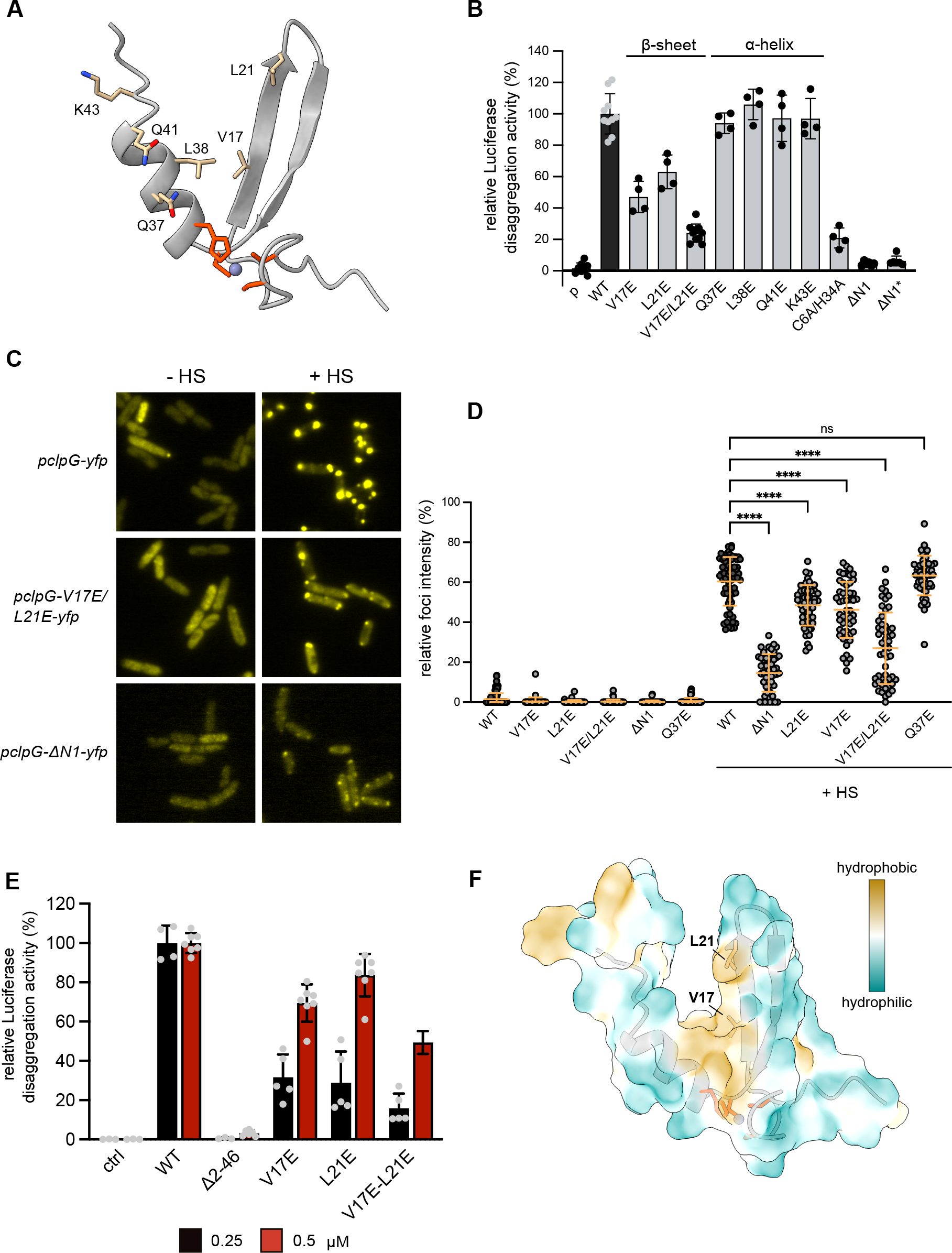
Val17 and Leu21 are crucial for ClpG targeting to protein aggregates. (A) Structure of the N1 core domain. Residues implicated in Zn^2+^ coordination are highlighted in red. Residues subjected to mutagenesis based on determined CSPs in presence of peptide 134 are indicated. (B) *E. coli t-*,.*c/pB* cells harboring plasmids for constitutive expression of Luciferase and IPTG-controlled expression of *clpG yfp* (wt and indicated mutants; vc: empty vector control) were grown at 30°C to mid-logarithmic growth phase. Cells were shifted to 46°C for 15 min and Luciferase activities were dete1mined during a recove1y phase at 30°C. The disaggregation activity of ClpG-YFP was set as 100%. (C) *E. coli t-*,.*c/pB* cells harboring plasmids for IPTG-controlled expression of *clpG-yfp* (wt and indicated mutants) were grown at 30°C to mid-log phase and shifted to 45°C for 15 min. Cellular localizations were dete1mined. (D) Distributions of diffuse and punctate YFP fluorescence intensities were detennined before and after heat shock (n=50). One-way Anova was applied to test for statistical significance. (E) Luciferase disaggregation activities of ClpG WT and indicated N1 mutants were detennined at indicated protein concentrations. A control reaction lacking ClpG is provided. (F) Hydrophobicity plot of the N1 core domain. Positions of Val1 7 and Leu21 residues are indicated. Standard deviations are based on at least three independent experiments (B, D/E).

We next explored whether the reduced disaggregation activity of ClpG N1 mutants correlates with a reduced binding to protein aggregates *in vivo*. ClpG-YFP largely displayed diffuse fluorescence at 30°C but massively relocalized to cell poles upon heat shock (Fig. 4C). Protein aggregates are deposited at the cell poles of *E. coli* cells and recruit disaggregating chaperones upon heat shock (30, 31). This indicates that stress-induced ClpG-YFP foci formation reflects binding to protein aggregates. The relative fraction of ClpG-YFP fluorescence intensity in stress-induced foci can therefore be used as readout for aggregate targeting. We determined this value for ClpG-YFP and N1 mutant variants that were most strongly affected in Luciferase disaggregation (Fig. 4D). While on average 60.4 % of ClpG-YFP fluorescence intensity was present in foci after heat shock, we observed less pronounced foci formation and increased diffuse fluorescence for disaggregation defective N1 mutants (Fig. 4C/D, Fig. S4B). The relative fluorescence intensity determined in foci was particularly low for ClpG-V17E/L21E (27% average foci intensity) and was similar to the aggregate-binding deficient controls ΔN1-ClpG-YFP lacking the N1 domain (14.6% average foci intensity) (Fig. 4C/D). ClpG-Q37E-YFP, which exhibited WT-like disaggregation activity, showed intense foci formation upon heat shock (63.4% average foci intensity) that was similar to ClpG-YFP. This demonstrates the specificity of the microscopic assay and underlines the correlation between reduced disaggregation activity and aggregate targeting. Heat resistance was tested to monitor the function of ClpG-V17E/L21E in *Pa* cells. ClpG-V17E/L21E was expressed at the same levels as ClpG WT, yet it exhibited reduced ability to restore heat resistance in a *Pa* Δ*clpB* Δ*clpG* Δ*clpG*_*GI*_ mutant, indicating reduced disaggregation activity (Fig. S4C/D).

We next characterized ClpG-V17E, L21E and V17E/L21E *in vitro*. We observed reduced disaggregation activities for all mutants, that became most obvious at low ClpG concentrations (Fig. 4E). The ClpG-V17E/L21E double mutant showed lowest disaggregation activity, consistent with our former *in vivo* findings. None of the mutants exhibited reduced ATPase activities excluding defects in the AAA+ threading motors as basis for disaggregation defects (Fig. S4E). The mutants were also not affected in Zn^2+^ binding (Fig. S4F), arguing against global unfolding of the N1 domain, which we confirmed by comparing 1D ^1^H-spectra of isolated N1* wild type and mutant derivatives (Fig. S4G). All mutant spectra exhibited the same peak dispersion as the wild type, which is a clear sign of a folded domain.

Together these results confirm the *in vivo* findings and substantiate the crucial role of V17 and L21 in aggregate binding and thus ClpG disaggregation activity. Both residues are part of a hydrophobic patch of the N1 core domain (Fig. 4F), which will enable N1 to interact with hydrophobic residues exposed on the surface of a protein aggregate.

### Multiple N1 domains are needed for efficient protein disaggregation

The estimated binding affinity of N1* to Peptide 134 is moderate (K_d_ > 50 μM). We therefore speculated that the presence of multiple N1 domains in a ClpG hexamer increases substrate affinity via avidity effects and asked how many N1 domains must be present in a hexamer to enable for efficient aggregate binding and disaggregation. We considered this point highly relevant as it mechanistically addresses ClpG substrate specificity: how can ClpG distinguish the surface of an insoluble protein aggregate from e.g. a soluble, non-native nascent polypeptide chain? We performed mixing experiments using N1*-ClpB-K476C and ΔN-ClpB-K476C as model system, since ClpB hexamers dynamically exchange subunits ensuring stochastic formation of mixed hexamers (32, 33). We confirmed mixing of WT and mutant subunits by showing that the presence of ATPase deficient ΔN-ClpB-E218A/K476C/E618A, harboring mutated Walker B motifs in both AAA domains, strongly poisoned disaggregation activity of N1*-ClpB-K476C (Fig. S5). Next, we determined the disaggregation activities of mixed N1*-ClpB-K476C/ΔN-ClpB-K476C hexamers generated at diverse mixing ratios. These experiments were done in a concentration-sensitive range as shown by linear correlation of N1*-ClpB-K476C concentrations and corresponding disaggregation activities (Fig. 5A). Addition of ΔN-ClpB-K476C to N1*-ClpB-K476C reduced disaggregation activities despite increasing total hexamer concentration, indicating poisoning upon incorporation of subunits that do not harbor N1* (Fig. 5A). To calculate the number of N1* required for protein disaggregation, we compared the relative disaggregation activities determined at diverse mixing ratios with those derived from a theoretical model, assuming that a mixed hexamer only displays activity if it contains a certain number of N1*-ClpB-K476C subunits (Fig. 5B). We found that approximately four N1* domains must be present in a hexamer to confer high disaggregation activity. In a reciprocal approach we tested whether an isolated N1* can inhibit ClpG or N1*-ClpB-K476C disaggregation activity by competing for aggregate binding (Fig. 5C). Addition of up to a 20-fold excess N1* did not reduce disaggregation activities. This suggests a vast increase in binding affinities of N1*-harboring AAA+ hexamers to protein aggregates and explains efficient outcompetition of isolated N1* by full-length ClpG harboring six N1 domains.

**Figure 5.**
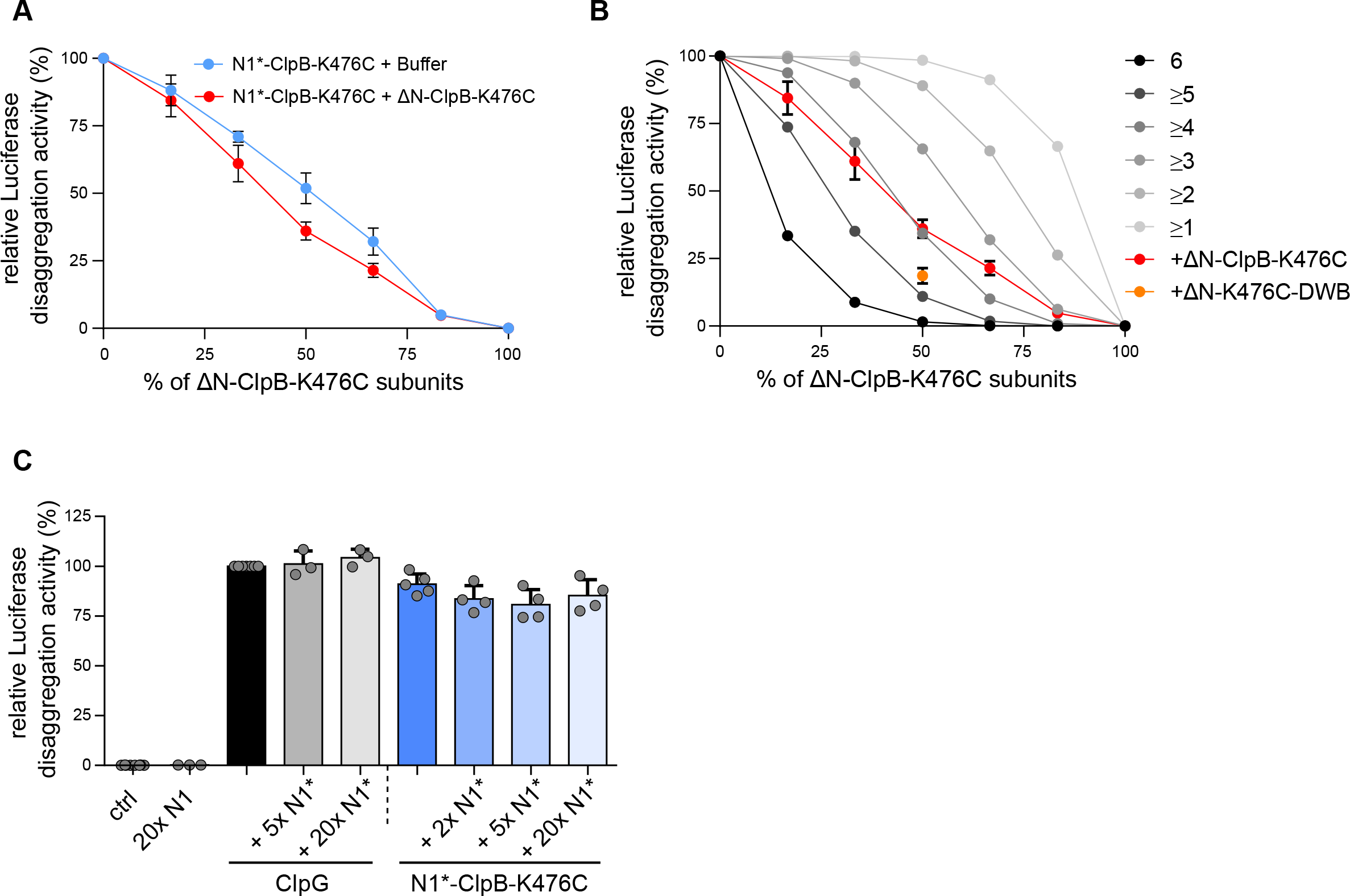
Multiple N1 domains are required for efficient disaggregation by ClpG hexamers. (A) Reactivation rates of aggregated Luciferase were determined in the presence of N1*-ClpB-K476C and AN-ClpB- K476C mixes and were set as 100% for non-mixed N1*-ClpB-K476C. Mixing ratios are indicated as % AN-ClpB-K476C. As reference, the disaggregation activities of lower N1*-ClpB-K476C concentrations (+ buffer) corresponding to the respective mixtures were determined. (B) Luciferase disaggregation activities of N1*-ClpB-K476C and ΔN-ClpB-K476C or ΔN-ClpB-DWB-K476C mixes were determined (red and orange, respectively). Disaggregation activities are compared with curves calculated from a model (black to grey), which assumes that a mixed hexamer only displays disaggregation activity if it contains the nunber of N1 domains indicated. Mixing ratios are indicated as % ΔN-ClpB-K476C. (C) Luciferase disaggregation activities of ClpG and N1*-ClpB-K476C were determined in absence and presence of isolated N1* as indicated. Disaggregation activity of ClpG inN1* absence was set to 100%. Standard deviations are based on at least three independent experiments (B-D).

## Discussion

Here, we report on the core structure of the aggregate targeting N1 domain of the autonomous ClpG disaggregase. The small domain is composed of an α-helix, an anti-parallel β-sheet and a Zn^2+^-coordinating center, which is crucial for N1 structural integrity. We identify V17 and L21 as conserved residues that play crucial roles for substrate binding. Both residues are part of a hydrophobic patch, which is well suited to interact with hydrophobic residues exposed on the surface of a protein aggregate. Notably, V17 and L21 are part of the first β-strand, while none of the mutations located in the N1* α-helix affected ClpG disaggregation activity. This might point to another feature of ClpG substrate selectivity next to hydrophobicity: the formation of β-sheets with β-strands of aggregated proteins. Such interaction is indeed predicted by AlphaFold2 for the model Peptide 134 (Fig. 2E). An increased β-sheet content of protein aggregates formed by heterologous proteins upon production in bacteria has been reported (34, 35). It is therefore tempting to speculate that the N1 domain interacts with unsaturated β-strands present on an aggregate surface via hydrophobic interactions. Notably, the N-terminal domains of the bacterial AAA+ protein ClpE and of the ClpC adaptor protein McsA exhibit sequence homologies with ClpG N1* and include a Zn^2+^-coordination center and hydrophobic residues at a position equivalent to N1*-V17 (Fig. S6A). Accordingly, AlphaFold2 predictions of ClpE and McsA NTD structures are similar to the ClpG N1* structure (Fig. S6B). *B. subtilis* ClpE and McsA have both been implicated in cellular protein disaggregation (36, 37). It is therefore tempting to speculate that the suggested functions of ClpE and McsA in protein disaggregation involve their homologous NTDs to target them to protein aggregates.

While hydrophobicity and, potentially, specific structural features of a substrate represent the basis for N1 interaction, a single binding event is not sufficient for recruitment of ClpG hexamers to protein aggregates. Instead, interactions between multiple N1 domains and various hydrophobic patches are required. We suggest that it is the presence of multiple hydrophobic patches located in close vicinity on an aggregate surface that represents the key recognition determinant. This feature enables for the simultaneous docking of multiple N1 domains creating strong avidity effects and directing the ClpG hexamer to protein aggregates. This binding mode provides strong substrate selectivity while preventing ClpG from targeting not yet completely folded nascent polypeptides and protecting those from ATP-fueled unfolding activities. This principle for aggregate recognition seems conserved since partner-dependent ClpB has to interact with multiple aggregate-bound Hsp70s for efficient aggregate recruitment and ATPase activation (16). Thus, while Hsp70 itself cannot discriminate between soluble non-native and aggregated proteins, it is the high local Hsp70 density on an aggregate surface that functions as specificity label for ClpB. This indirect recognition principle has been changed to a direct one in case of the autonomous ClpG disaggregase. We also envision that multiple anchoring points between ClpG and the aggregate surface enhance pulling forces, locally loosening contacts between unfolded proteins within the aggregate and facilitating initialization of the threading process.

Next to its role as substrate binding platform, we show that N1 additionally functions as negative regulator of ClpG, as all N1 deletion constructs and the Zn^2+^-binding deficient mutant C6A/H34A exhibited an enhanced ATPase activity. Accordingly, ClpG-Δ47-71, which is proficient in aggregate binding, functioned as superior disaggregase as compared to ClpG WT. How N1 downregulates ClpG activity remains to be determined. The dual role of N1 in substrate binding and ClpG activity control opens a pathway to activate the disaggregase on demand. For future studies, it will be now important to dissect how N1 represses ClpG activity and how substrate binding triggers activation.

### Experimental procedures

#### Strains and plasmids

All strains and plasmids used in this study are summarized in Table S1. *Escherichia coli* cells were grown in LB medium at 30°C containing appropriate antibiotics with agitating speed 120 rpm. *E. coli* XL1 blue was used for cloning and retaining of plasmids requiring kanamycin (Km) at 50 μg ml^−1^ and ampicillin (Ap) at 100 μg ml^−1^ for plasmid propagation.

### Protein purification

*E. coli* ClpB (wt and derivatives) was purified after overproduction from *E. coli* Δ*clpB::kan* cells using pDS56-derived expression vectors (27). *P. aeruginosa* ClpG_GI_ (wild type and mutant derivatives) and N1*-ClpB fusion constructs were purified after overproduction in *E. coli* BL21 cells using pET24a-derived expression vectors. ClpG_GI_ deletion mutants and N1*-ClpB hybrids were generated by PCR and point mutants were constructed by quickchange one step site directed mutagenesis. All mutations were verified by sequencing. All proteins harbor a C-terminal His_6_-tag and were purified using Ni-IDA (Macherey-Nagel) following the instructions provided by the manufacturer. In short, cell pellets were resuspended in buffer A (50 mM NaH_2_PO_4_, 300 mM NaCl, 5 mM β-mercaptoethanol, pH 8.0) supplemented with protease inhibitors (Roche). After cell lysis using French press the cell debris was removed by centrifugation at 16000 g for 45 min at 4°C and the cleared lysate was incubated with Protino IDA resin (Macherey-Nagel) for 1 h at 4°C. Afterwards the resin was transferred into a plastic column and washed once with buffer A. His-tagged proteins were eluted by addition of buffer A supplemented with 250 mM Imidazole. Subsequently, pooled protein fractions were subjected to size exclusion chromatography (Superdex S200, Amersham) in buffer A1 (50 mM Tris pH7.5, 50 mM KCl, 10 mM MgCl_2_, 5% (v/v) glycerol, 2 mM DTT) for ClpB and buffer A2 (50 mM Tris pH7.5, 50 mM KCl, 20 mM MgCl2, 5% (v/v) glycerol, 2 mM DTT) for ClpG.

For NMR experiment Strep-tagged N1* was produced in *E. coli* BL21 cells in M9 medium with 1g/l ^15^NH_4_Cl and 2 g/l ^13^C-glucose. The domain was purified using Strep-Tactin Superflow High capacity resin (IBA-lifescience) equilibrated in Strep-Tactin buffer (100 mM Tris pH 8.0, 150 mM NaCl, 5 mM β-mercaptoethanol). Bound proteins were eluted using Strep-Tactin buffer supplemented with 12.5 mM d-desthibiotin (Merck). Fractions with highest protein content were pooled and subjected to SEC using a HiLoad 16/600 Superdex S30 pg (GE Healthcare) column equilibrated in 50 mM Tris pH 7.5, 50 mM KCl, 20 mM MgCl_2_, 2 mM DTT. Fractions with highest protein content were pooled and concentrated using Centricons (3 – 5 kDa cutoff) pre-equilibrated with NMR buffer (50 mM NaH_2_PO_4_ pH 6.5, 50 mM KCl, 2 mM DTT).

Purifications of DnaK, DnaJ, GrpE and Firefly Luciferase were performed as described previously (16, 27, 32). Pyruvate kinase of rabbit muscle and Malate Dehydrogenase of pig heart muscle were purchased from Sigma. Protein concentrations were determined with the Bradford assay (Biorad).

### *In vitro* and *in vivo* disaggregation assays

200 nM Firefly Luciferase were heat-aggregated at 47°C for 30 min or 46°C for 15 min respectively in assay buffer (50 mM Tris pH 7.5, 50 mM KCl, 20 mM MgCl_2_, 2 mM DTT). Aggregated proteins were mixed (final concentration 100 nM) 1:1 with disaggregases (final concentrations: 0.25-1 μM ClpG (wild-type or mutants or 1 μM ClpB (wild type or mutants) with 1 μM DnaK, 0.2 μM DnaJ, 0.1 μM GrpE (KJE)). For N1-ClpB-K476C mixing experiments the final chaperone concentration was 150 nM. The disaggregation reaction was started by addition of an ATP regenerating system (2 mM ATP, 12 mM Phosphoenolpyruvate, 20 ng/μl Pyruvate kinase) and performed at 30°C in assay buffer. Luciferase activities were determined using a Lumat LB 9507 or a Lumat LB 9510 (both Berthold Technologies). 2 μl of disaggregation reaction was mixed with 100 μl luciferase assay buffer (25 mM glycylglycine pH 7.4, 12.5 mM MgSO_4_, 5 mM ATP) and subsequently 100 μl luciferin (Gold Biotech) were injected. 100% activity corresponds to Luciferase activity before heat denaturation.

For *in vivo* luciferase disaggregation *E. coli ΔclpB* cells harboring *placI*^*q*^*-luciferase* and either *pUHE21-clpG* derivatives (wt and mutants) or *pDK66-clpG*-*yfp* derivatives (wt and mutants) were grown in LB medium supplemented with ampicillin (100 μg/ml) and spectinomycin (50 μg/ml) at 30 °C while shaking at 120 rpm to early-logarithmic phase. *clpG*/*clpG-yfp* expression was induced by addition of IPTG (25 μM, for *clpG-*Δ*N1*: 1000 μM) for 2 h. Production of chaperones to similar levels was verified by SDS-PAGE and subsequent Coomassie staining. After 2 h (mid-logarithmic phase), native Luciferase levels were determined in a Lumat LB 9507 and set to 100%. For that, 100 μl of cells were transferred into plastic tubes, 100 μl of 250 nM Luciferin were injected und luminescence was measured for 10 s. Next, 900 μl aliquots of cells were shifted to 46 °C for 15 min to induce a non-lethal heat-shock. Immediately afterwards, tetracycline (70 μg/ml) was added to stop protein synthesis and cells were moved back to 30 °C. Luciferase activities were determined after 0, 15, 30, 60, 90 and 120 min during the recovery phase and the efficiency of the disaggregation reaction was determined at respective timepoints.

### ATPase Assay

The ATPase activities of 0.25-1 μM ClpG wild-type or mutants were determined in a reaction volume of 100 μl in assay buffer with 0.5 mM NADH (Sigma), 1 mM PEP (Sigma) and 1/100 (v/v) PK/LDH mix (Sigma). 100 μl of 4 mM ATP in assay buffer (50 mM Tris pH 7.5, 50 mM KCl, 20 mM MgCl_2_, 2 mM DTT) was added to each reaction in a 96-well plate (TPP) format to start the reaction. The decrease of NADH absorbance at 340 nm was determined in a BMG Labtech Clariostar Omega plate reader at 30°C. ATPase activities were calculated by assuming a 1:1 stoichiometry of NAD^+^ oxidation and the production of ADP.

### Heat Resistance Assay

*E. coli ΔclpB* cells harboring pUHE21 derivates allowing for IPTG controlled expression of *clpG* (wild-type or mutants) were grown in LB media at 30°C to early logarithmic phase (OD_600_: 0.15 – 0.2). Expression of the respective proteins was induced by addition of 100 μM IPTG. Protein production was documented 2 h after IPTG addition by western blot analysis. Subsequently one ml aliquots were shifted to 50°C for 120 min. At indicated time points bacterial survival was determined by preparing serial dilutions, spotting them on LB plates followed by incubation for 24 h at 30°C. *P*.*aeruginosa* SG17M Δ*clpB* Δ*clpG* Δ*clpG*_*GI*_ cells harboring pJN105 derivates allowing for L-arabinose controlled expression of *clpG* (wild-type or mutants) were grown in LB media at 37°C to early logarithmic phase (OD_600_ = 0.2 – 0.4). 1% of L-arabinose was used to express the respective proteins. After incubation at 37°C until OD_600_ = 0.8, cell suspensions were exposed to 50°C for 30 and 60 min. At indicated time points bacterial survival was determined by preparing serial dilutions and subsequent spotting on LB plates followed by incubation for 18 h at 37°C.

### CD spectroscopy

CD spectra of 20 μM N1 1-46 (wt or Zn^2+^ binding mutant) in 10 mM Na phosphate buffer pH 7.5 (pH 7.5) were recorded using a Jasco J750 spectropolarimeter.

### Size exclusion chromatography

Oligomerization of N1 domains was monitored by size exclusion chromatography (SEC) using a Superdex S30 10/300 column (GE Healthcare) at 4°C. The column was equilibrated with assay buffer 30 μM N1 domain was incubated for 5 min at room temperature before injection. Elution fractions were analysed by SDS-PAGE followed by Sypro staining. Carbonic anhydrase (29 kDa), Rnase A (13.7 kDa) and Aprotinin (6.5 kDa) served as molecular mass standards (all GE Healthcare). Sypro Ruby Protein Gel Stain (Molecular probes) was performed as instructed by the manufacturer.

### Bioinformatic analysis

Multiple sequence alignments were performed using Clustal Omega (https://www.ebi.ac.uk/Tools/msa/clustalo/) and were displayed using Jalview.

### Determination of Zn^2+^ binding by ICP-OES

Zn^2+^-binding of ClpG wild-type and mutants was determined via Inductively Coupled Plasma Optical Emission Spectrometry (ICP-EOS). 0.9 ml of 10 mM proteins (20 mM Tris-HCl pH 7.5, 10 mM KCl, 5 % glycerol, 2 mM DTT) were incubated with 2 ml HNO_3_ (65 %, p.A., NeoFroxx GmbH) for 1 h at 90°C. Afterwards double-distilled water was added to a final volume of 10 ml. Zn^2+^ presence was determined using Agilent 720 ICP-OES at a wavelength of 213,857 nm. For calibration ICP multi-element-standard-solution IV (Merck KGaA) was used.

### Fluorescence Microscopy

*E. coli ΔclpB* cells harboring *placI*^*q*^*-luciferase* and *pDK66-clpG*-*yfp* derivatives (wild-type and mutants) were grown in LB medium supplemented with ampicillin (100 μg/ml) and spectinomycin (50 μg/ml) at 30°C while shaking at 120 rpm to early-logarithmic phase. *clpG-yfp* expression was induced by addition of IPTG (25 μM, for *clpG-*Δ*N1*: 1 mM) for 2 h. Production of chaperones to similar levels was verified by SDS-PAGE and subsequent Coomassie staining. After 1 h 45 min, 1 ml of cells was centrifuged at 4000 rpm for 1 min at room temperature. 850 μl of supernatant was removed and the remaining cell pellet was resuspended in the remaining volume. 2 μl of the suspension were applied to a cover slip (22 × 60 mm, Menzel) and the cells were mechanically fixed with agarose pads (1% (w/v) in 0.5x TBE). Images were acquired using an Olympus CellSens IX81 inverted microscope using a PLAPO 100x/1.45 Oil DIC immersion objective using YFP filters. Cells were heat-shocked at 45 °C for 15 min in a water bath while shaking at 120 rpm and images were taken immediately afterwards as described above. Image analysis for determination of the foci intensity was performed using ImageJ.

### Western Blotting

Total extracts of cells were prepared and separated by SDS-PAGEs, which were subsequently electrotransferred onto a PVDF membrane. The membrane was incubated in blocking solution (3% BSA (w/v) in TBS) for 1 h at RT. Protein levels were determined by incubating the membrane with ClpG_GI_-specific antibodies (1:10.000 in TBS-T + 3% (w/v) BSA) and an anti-rabbit alkaline phosphatase conjugate (Vector Laboratories) as secondary antibody (1:20.000). Blots were developed using ECF™ Substrate (GE Healthcare) as reagent and imaged via Image-Reader LAS-4000 (Fujifilm).

### Nuclear magnetic resonance spectroscopy

All NMR experiments were recorded at 298 K using a Bruker Avance III 700 MHz spectrometer equipped with a triple-resonance probe. Protein concentrations were in the range between 80 and 800 μM. Assignments were obtained using standard ^1^H,^13^C,^15^N scalar correlation experiments (38) employing apodization weighted sampling (39). The data were processed with NMRpipe (40) and analysed with NMRView (41). NOE-based distance restraints were obtained from ^15^N- and ^13^C-edited 3D NOESY-HSQC, and ^13^C-edited 3D HMQC-NOESY experiments. NOE assignments were performed manually using NMRView. Structures were calculated in the ARIA1.2/CNS1.2 package (42) with standard simulated annealing protocols using a log harmonic NOE potential for the distance restraints and a patch for the coordination of the Zn^2+^ ion (29). Dihedral angle restraints obtained from TALOS+ were also included in the structure calculation (28).

^15^N spin relaxation parameters *R*_*1*_, *R*_*2*_, and heteronuclear NOEs were recorded on the same spectrometer and conditions. For *R*_*1*_ experiments, relaxation delays of 20, 50, 100, 150, 250, 400, 500, 650, 800, 1000, 1300 and 1600 ms were used, where the 150 ms delay was acquired in duplicates to estimate peak volume uncertainties. For *R*_*2*_ experiments, relaxation delays of 16, 32, 48, 64, 80, 96, 112, 128, 160, and 192 ms were used, where the 16 and 80 ms delays were acquired in duplicates to estimate peak volume uncertainties. To derive the order parameter *S*^*2*^ from Lipari-Szabo model-free analysis (43, 44) we employed the program ROTDIF (45). The chemical shift perturbations (CSPs) plotted in Figure XX were calculated according to CSP = {0.5*((0.14ΔdN)2 + (ΔdH)2)}1/2, where ΔdN and ΔdH are the chemical shift changes of ^15^N and ^1^HN, respectively, comparing free protein N1* (80 μM) and in the presence of peptide 134 (240 μM).

## Data availability

All data are contained within the manuscript. The NMR structure ensemble of N1* has been deposited at the PDB (PDB ID 8P66).

## Acknowledgments

P.K. and T.J. were supported by the Heidelberg Biosciences International Graduate School (HBIGS). We thank Christian Scholz (Institut für Geowissenschaften, University of Heidelberg) for performing ICP-EOS measurements. Janosch Hennig gratefully acknowledges support from the European Molecular Biology Laboratory.

## Author contributions

Conceptualization, P.K. J.H. and A.M.

Methodology, P.K., B.S., J.H. and A.M.

Investigation, P.K.,B.S., S.M., C.L., J.H. and A.M.

Formal Analysis, P.K., B.S., T.J.,S.M., C.L., J.H. and A.M.

Resources, P.K. and A.M.

Writing – Original Draft, A.M.

Writing – Review and Editing, P.K., B.S., T.J. C.L., J.H.

Supervision, A.M.

Visualization, P.K. and A.M.

Funding Acquisition, C.L., A.M.

## Funding and additional information

This work was supported by a grant of the Deutsche Forschungsgemeinschaft (MO970/7-1) to A.M. C.L. received funding from the National Research Foundation of Korea (NRF) funded by the Korea government (MSIT) (grant 2021R1C1C1011690), the Core Research Institute Basic Science Research Program through the NRF funded by the Ministry of Education (grant 2021R1A6A1A10044950), and the new faculty research fund of Ajou University.

