## Supplementary Figures 1-6 for "Structural basis of aggregate binding by the AAA+ disaggregase ClpG"

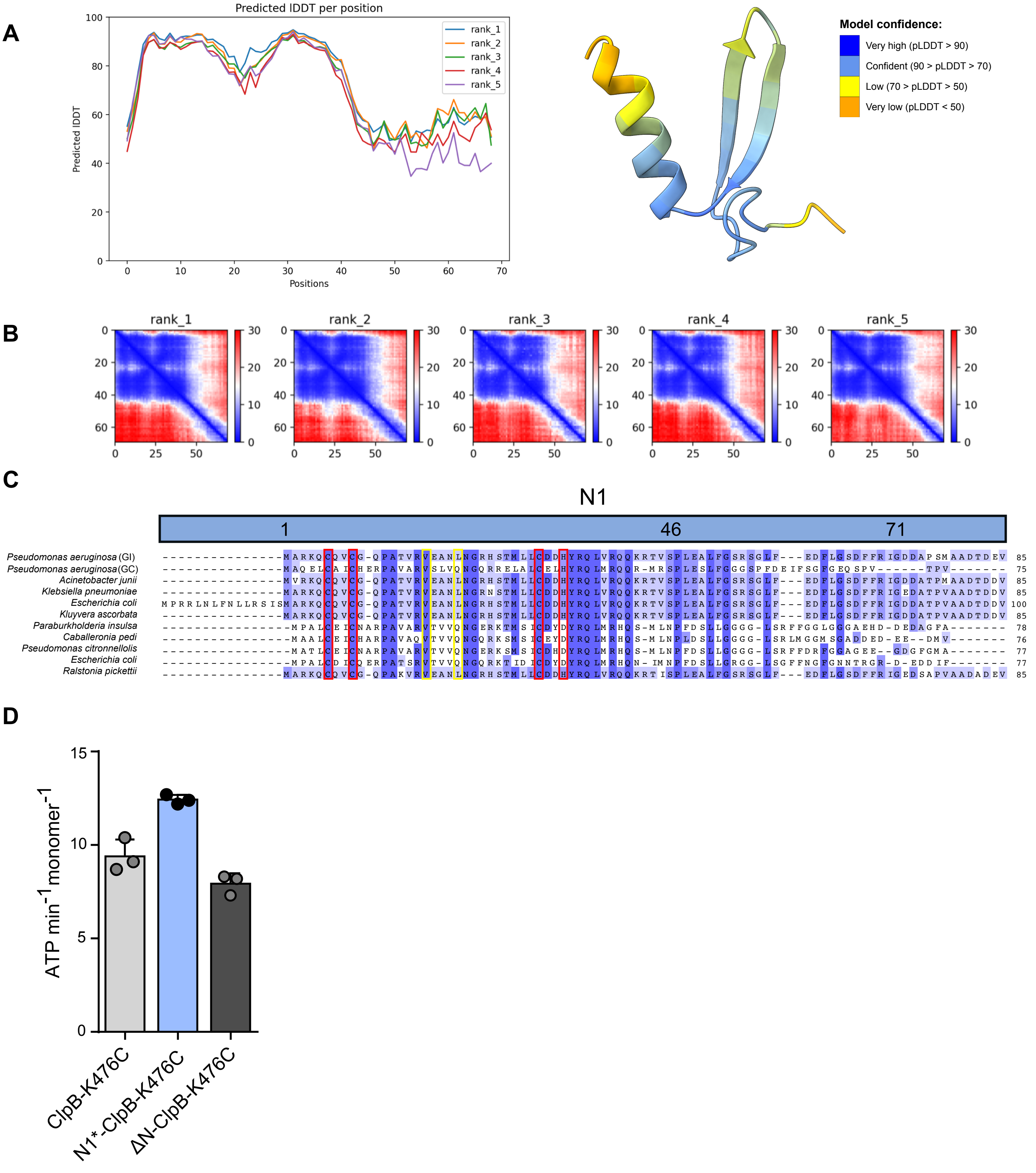


**Figure S1**

AlphaFold2 structure prediction and sequence alignment of the ClpG N1 domain. (A) pLDDT values for five N1 domain structure predictions indicating that the core domain, residues 5-40, are predicted with high confidence, whereas the C-terminal region, although predicted to be helical, has a low confidence. The structural model on the right depicts the calculated confidence with the indicated colour code. (B) Predicted alignment error for five N1 domain structure predictions indicating similarly high likelihoods for the core domain. (C) Sequence alignment of the N1 domains of indicated ClpG homologs. Boundaries of N1 subdomains are indicated. Similar and identical residues are highlighted in light and dark blue. Residues implicated in binding to $Zn^{2+}$ and protein aggregates are highlighted in red and yellow, respectively. (D) ATPase activities of ClpB-K476C, N1*-ClpB-K476C and ΔN-ClpB-K476C were determined. Standard deviations are based on at least three independent experiments (D).

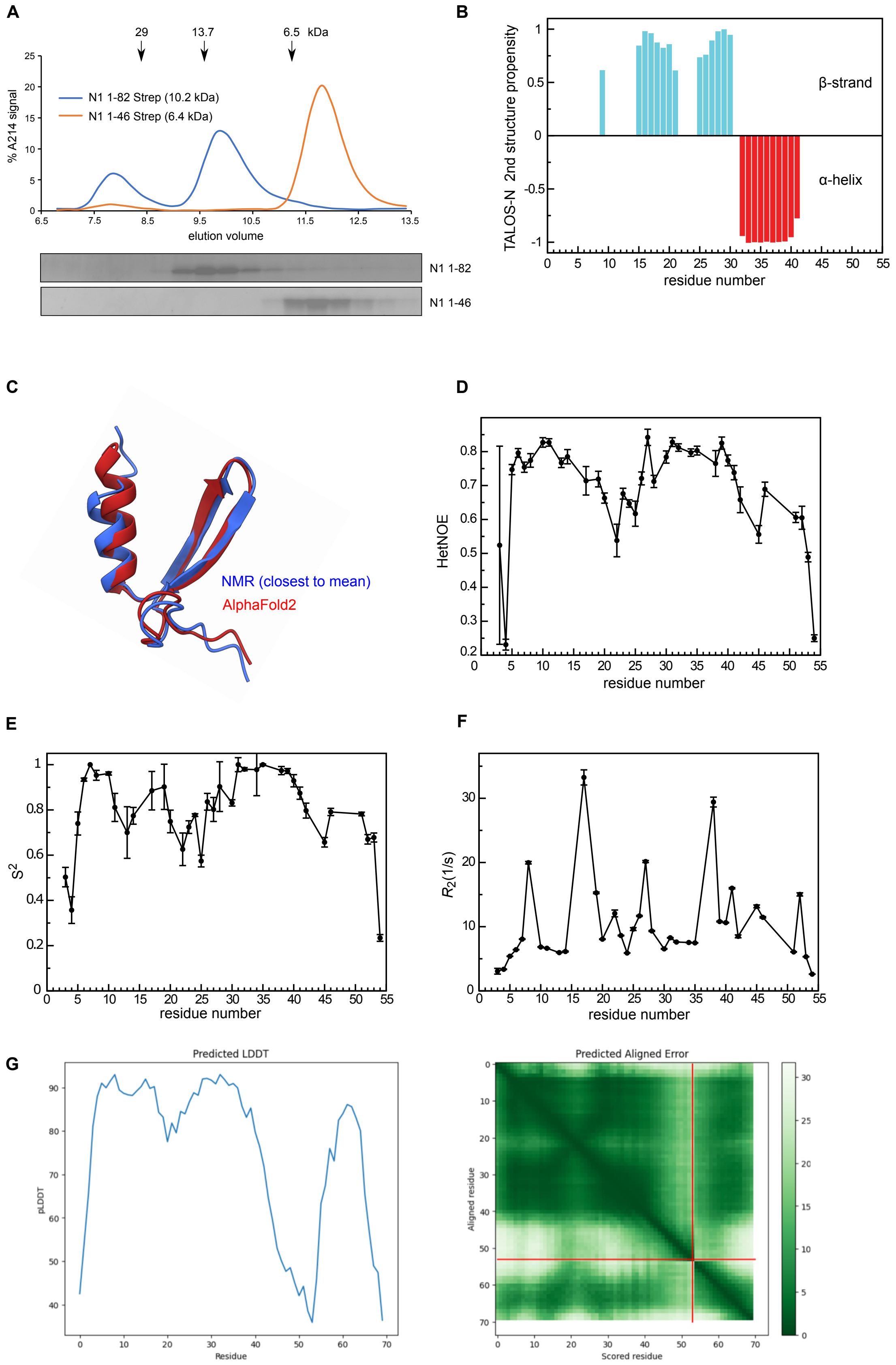


### Figure S2

Characterization of the N1 domain by SEC and NMR. (A) N1 domains (1-46, 1-82) harboring a C-terminal Strep-tag were subjected to S30 size exclusion chromatography. Elution profiles were recorded and respective fractions analyzed by SDS-PAGE and Sypro Ruby staining. Elution positions of marker proteins are provided. (B) Secondary structure propensity predicted from secondary chemical shifts using TALOS. The predicted structure is in accordance with the AlphaFold2 model and the NMR structure of the N1* domain. (C) Superposition of the AlphaFold2 prediction of the N1* domain (red) and the NMR structure closest to the mean (blue). (D) Heteronuclear NOE values per residue indicate higher flexibility in the C-terminal region and around residues 17-27. (E) Order parameter $S^2$ per residue, derived from Lipari-Szabo model-free analysis indicate that the region around residues 13-27 and 42-54 are dynamic. (F) Measured transverse relaxation rates ($R_2$) indicate that V17 (exhibiting the strongest CSPs upon titration with peptide 134), T27 and L38 change conformation in the slow exchange regime. (G) The pLDDT and pAE plots to assess the AlphaFold2 prediction confidence of the N1*-peptide 134 complex are shown.

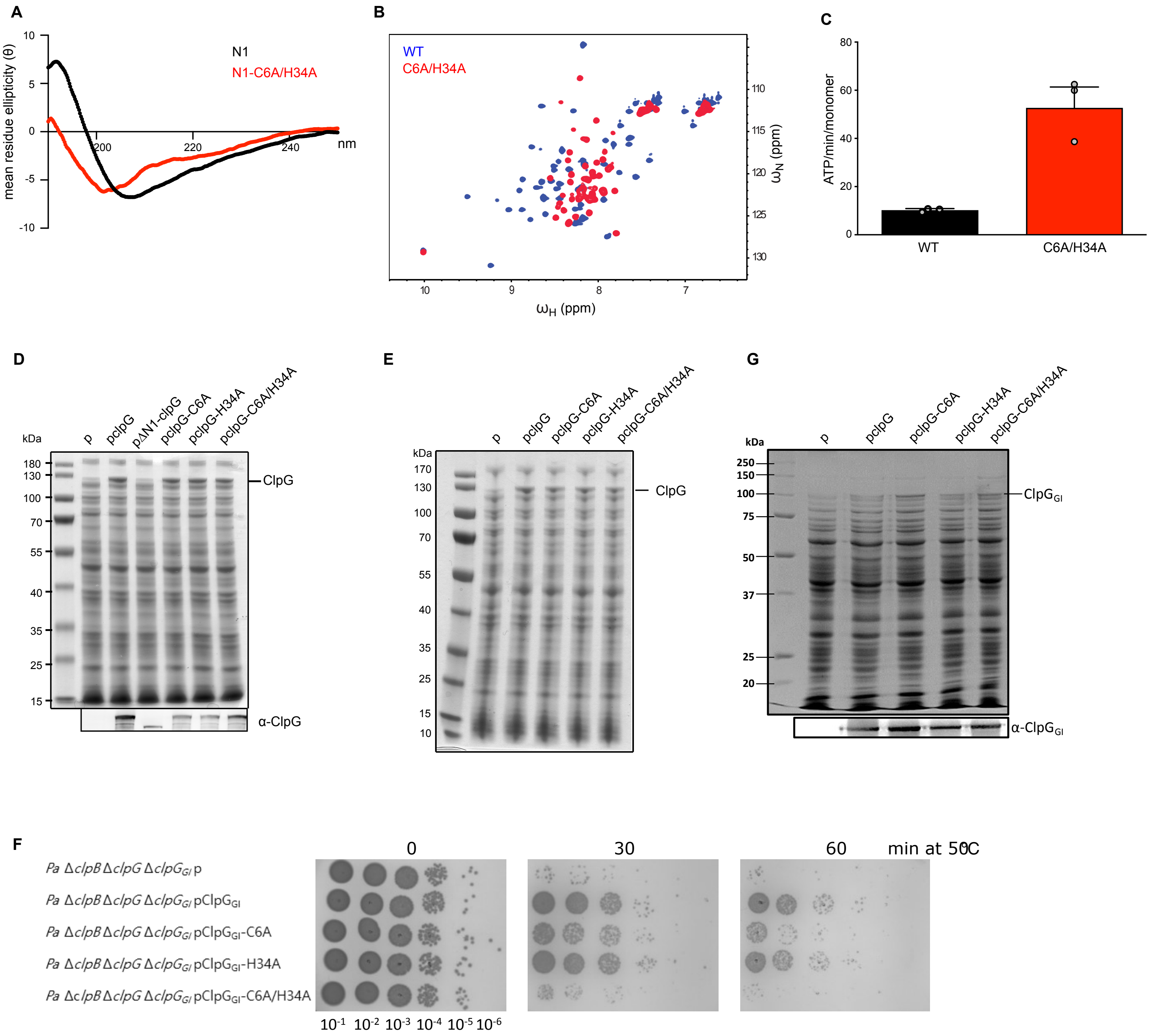


**Figure S3**

Characterization of the Zn$^{2+}$-binding mutant ClpG-C6A/H34A. (A) CD spectra of N1* and N1*-C6A/H34A. (B) Overlay of $^{1}$H-$^{15}$N HSQC spectra of N1* WT (blue) and N1*-C6A/H34A (red) show that resonances move from being well-dispersed in the WT to the middle of the spectrum for the mutant. This indicates that the double mutant C6A/H34A is unfolded. (C) ATPase activities of ClpG WT and ClpG-C6A/H34A were determined. (D-E) Production levels of ClpG (WT and mutants) in *E. coli ΔclpB* cells harboring *placI$^q$*-Luciferase (D) or *placI$^q$* (E). Cells were were grown at 30°C for 1.5 h. Expression of *clpG* (wild type or mutants) was induced from the pUHE21 expression vector by addition of 100 µM IPTG for 2 h. Total cell extracts were prepared and levels of ClpG were determined SDS-PAGE followed by Coomassie-staining. ClpG levels were additionally determined by western blot analysis using ClpG-specific antibodies (D). *p*: empty vector control. Standard deviations are based on at least three independent experiments (C). (F/G) *P. aeruginosa* (*Pa*) SG17M Δ*clpB* Δ*clpG$_{core}$* Δ*clpG$_{GI}$* cells harboring plasmids for expression of ClpG (WT and mutants) were grown at 37℃ to mid-logarithmic growth phase and shifted to 50℃. Serial dilutions of cells were prepared at the indicated time points, spotted on LB plates and incubated at 37℃ (F). Expression levels were determined prior to heat shock by SDS-PAGE and similar ClpG production levels were confirmed by Coomassie staining and western blot analysis. *p*: empty vector control.

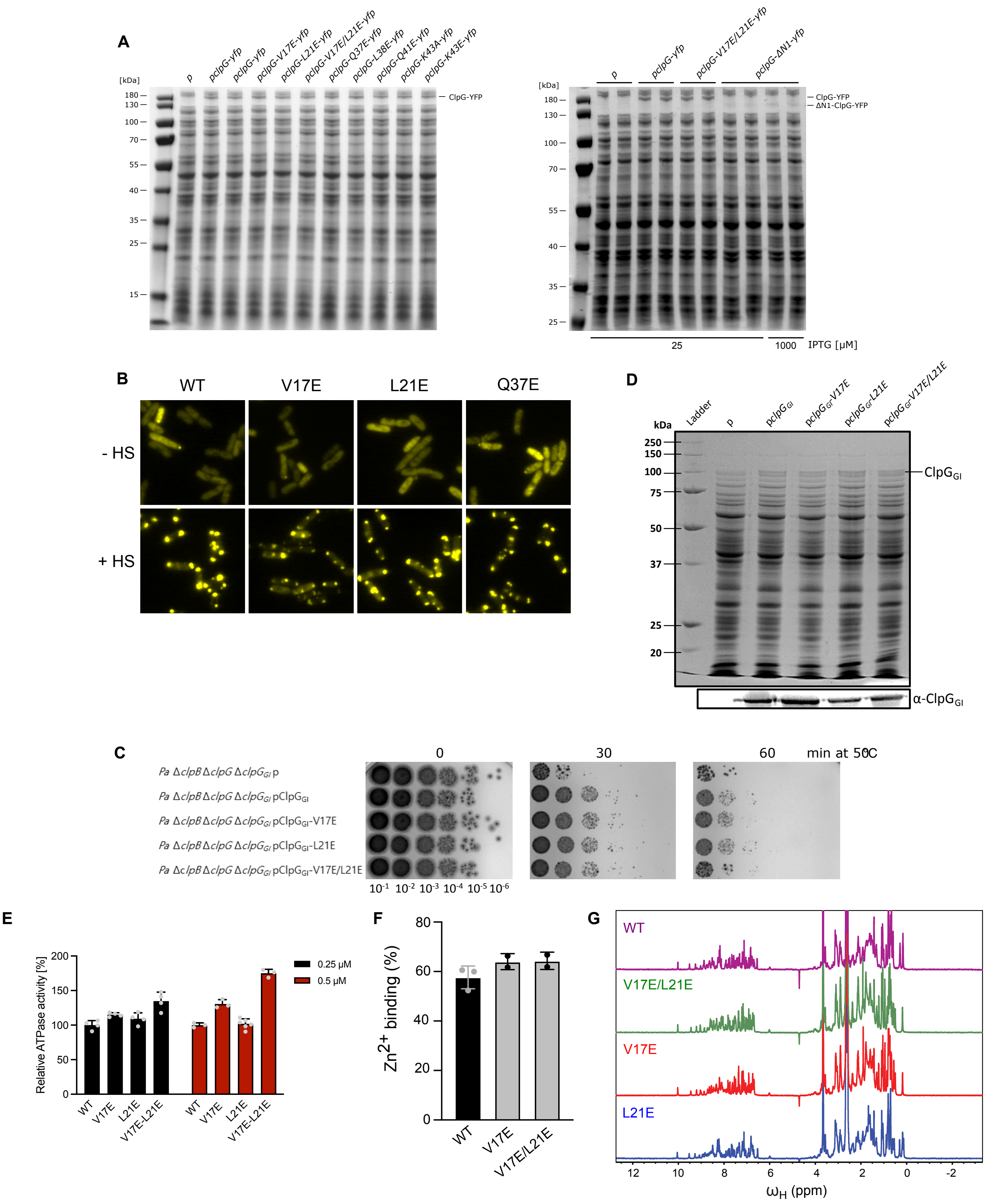


**Figure S4**

Analysis of N1 mutants. (A) Production levels of ClpG-YFP (WT and mutants) in *E. coli ΔclpB* cells harboring *placI^q*-Luciferase. Cells were were grown at 30°C for 1.5 h. Expression of *clpG* (wild type or mutants) was induced from the pDK66 expression vector by addition of 25 μM IPTG or 1 mM IPTG (ΔN1-ClpG-YFP) for 2 h. Total cell extracts were prepared and levels of ClpG-YFP were determined via SDS-PAGE followed by Coomassie-staining. *p*: empty vector control. (B) *E. coli ΔclpB* cells harboring plasmids for IPTG-controlled expression of *clpG-yfp* (wt and indicated mutants) were grown at 30°C to mid-log phase and shifted to 45°C for 15 min. Cellular localizations were determined. (C/D) *P. aeruginosa* (*Pa*) SG17M Δ*clpB* Δ*clpG$_{core}$* Δ*clpG$_{GI}$* cells harboring plasmids for expression of ClpG (WT and mutants) were grown at 37℃ to mid-logarithmic growth phase and shifted to 50℃. Serial dilutions of cells were prepared at the indicated time points, spotted on LB plates and incubated at 37℃ (C). Expression levels were determined prior to heat shock by SDS-PAGE and similar ClpG production levels were confirmed via Coomassie staining and western blot analysis (D). *p*: empty vector control. (E) ATPase activities of indicated ClpG variants were determined at 0.25 and 0.5 μM protein concentrations. ATPase activities of WT were set as 100%. (F) $Zn^{2+}$-binding of ClpG and indicated mutants was determined by ICP-OES. (G) Stacked $^{1}$H-1D NMR spectra of N1* and indicated mutant derivatives. The similar degree of peak dispersion in the amide and methyl region indicates that all mutants are folded.

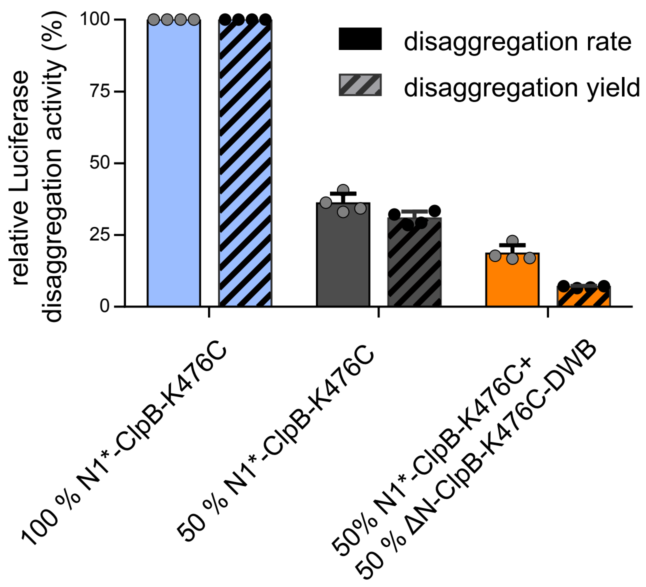


**Figure S5**

Disaggregation rates and yields of N1*-ClpB-K476C/ΔN-ClpB-K476C-DWB mixes. Disaggregation activities of 0.15 µM N1*-ClpB-K476C (100%), 0.075 µM N1*-ClpB-K476C (50%) and a mixture of 0.075 µM N1*-ClpB-K476C and 0.075 µM N1*-ClpB-K476C-DWB (ATPase-deficient). Refolding rates (% refolded Luciferase/min) and yields (% refolded Luciferase after 120 min) were determined and set to 100% for 0.15 µM N1*-ClpB-K476C. Standard deviations are based on at least three independent experiments.

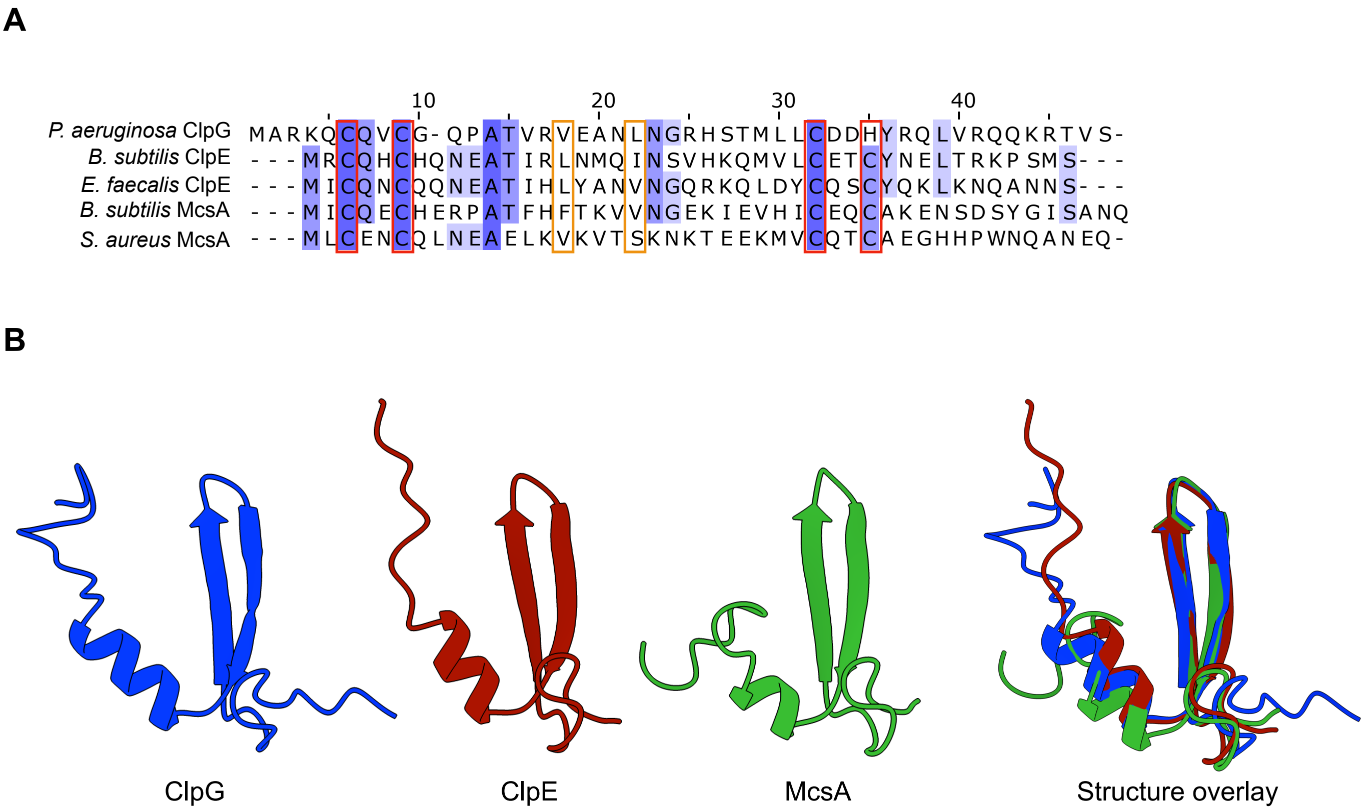


**Figure S6**

Sequence alignment of the *P. aeruginosa* ClpG N1* domain and N-terminal domains of *B. subtilis/E. faecalis* ClpE and *B. subtilis/S. aureus* McsA (A). Similar and identical residues are highlighted in light and dark blue. Residues implicated in binding to $Zn^{2+}$ and protein aggregates are highlighted in red and orange, respectively. (B) Comparison of the mean NMR structure of *P. aeruginosa* ClpG N1* with AlphaFold2 predictions of the N-terminal domains of *B. subtilis* ClpE and McsA.
